# AKAP12 variant 1 knockout enhances vascular endothelial cell motility

**DOI:** 10.1101/2025.02.02.635915

**Authors:** Ashrifa Ali, Bhaskar Roy, Micah B. Schott, Bryon D. Grove

## Abstract

In this study, we examined the role of AKAP12, in endothelial cell motility, with a specific focus on AKAP12 variants AKAP12v1 and AKAP12v2. Previous work has shown that AKAP12, a multivalent A-kinase anchoring protein that binds to PKA and several other proteins regulating protein phosphorylation, is expressed at low levels in most endothelia *in vivo* but at higher levels in cells *in vitro*. Here, we found that AKAP12 expression in endothelial cell (HUVEC) cultures was cell density-dependent, with the expression being highest in subconfluent cultures and lowest in confluent cultures. AKAP12 expression was also elevated in cells at the wound edge of wounded endothelial cell monolayers. Knockdown of variants 1 and 2 inhibited cell migration, whereas CRISPR/Cas9 knockout of AKAP12v1 enhanced migration, indicating that the absence of this variant and the presence of AKAP12v2 may shift the signaling pathways. Further analysis using bulk RNA sequencing revealed that the loss of AKAP12v1 affects genes associated with cell migration and intercellular junctions. We propose that AKAP12v1 and AKAP12v2 work together to modulate endothelial cell migration, providing insights into their distinct yet complementary roles in endothelial function and potential implications for cardiovascular health.

## Introduction

The vascular endothelium regulates a wide variety of physiological functions, including the maintenance of vascular tone through the regulation of vasodilation and constriction, molecular transport, cellular trafficking across the vascular wall, thrombogenesis, inflammation, and angiogenesis. Dysregulation within the endothelium significantly affects these physiological functions and, as a result, is considered a hallmark of cardiovascular disease [1] and an underlying factor in diseases such as stroke, heart disease, diabetes, insulin resistance, tumor growth, metastasis, and viral infectious disease [2–8]. Hence, understanding the regulation of endothelial cell processes, such as cell migration, proliferation, and maintenance of barrier function, is critical to understanding normal vascular function and the causes of vascular disease.

Endothelial cell function is controlled by a host of external factors that regulate cellular outcomes, such as proliferation, migration, and maintenance of barrier function, through a complex network of interconnected signaling pathways. The integration and regulation of this complex signaling network remain poorly understood, but despite this, it is now clear that scaffolding proteins such as A-Kinase Anchoring Proteins (AKAPs), PKC scaffolds RACK1 and annexins, and MAPK scaffold KSR [9–22] provide a molecular framework for directing signaling proteins toward specific substrates to favor one signal transduction pathway over another. Among these, AKAP12 has gained traction as a critical regulator of cellular functions in various cell types, including endothelial cells.

Human AKAP12 (also termed Gravin) was first identified in endothelial cells during screening of a human endothelial cell library with serum from a patient with myasthenia gravis [23]. Subsequent studies have revealed that AKAP12 is widely expressed in vertebrate cells and binds to a number of signaling proteins, including PKA, PKC, Src, and Cyclin D [23–35]. In addition, AKAP12 localizes to the plasma membrane due to the presence of several membrane localization domains, but translocates to intracellular locations in response to either activation of PKC or increased intracellular calcium concentration, resulting in changes in signaling protein localization [34, 36–39]. Furthermore, AKAP12 is expressed as two variants in endothelial cells, AKAP12v1 and AKAP12v2, controlled by individual promoters [40]. They are 95% identical, but differ in their N-terminal sequence, with AKAP12v1 containing an N-terminal myristoylation site and AKAP12v2 lacking it.

AKAP12 is normally expressed at low levels in endothelial cells *in vivo*, but is highly expressed in cultured endothelial cells, indicating that it may be upregulated under conditions that promote migration and cell proliferation [23, 28, 29]. AKAP12 knockdown in endothelial cell monolayers using siRNA impairs the cAMP-dependent increase in vascular endothelial barrier function, while overexpression of AKAP12 has been shown to promote barrier function [41, 42], implying that AKAP12 is likely important in regulating vascular integrity and permeability. However, the roles of the two AKAP12 variants in endothelial cell migration and barrier formation are unknown. Studies on the effect of AKAP12 expression on cell migration, proliferation, and angiogenesis have, for the most part, either not considered the role of each AKAP12 variant or overexpressed AKAP12v1 using expression vectors. To date, the only study focused on AKAP12v2 was a study by Finger et al. [43], who reported that hypoxia specifically increased AKAP12v2 in melanoma cells, contributing to changes in PKA phosphorylation profiles at the membrane and increased metastasis. Except for this study, the role of individual variants of AKAP12 has not been comprehensively investigated and has never been studied in vascular endothelial cells.

In this study, we examined the role of AKAP12 and its variants in endothelial cell migration *in vitro*. We first demonstrated that the expression of AKAP12 is regulated in a cell density-dependent manner, with AKAP12 expression being lowest in confluent monolayers and highest in low-density cells. In addition, wounding of monolayers resulted in increased AKAP12 expression at the wound edge in wounded monolayers. Knockdown of both AKAP12 variants with antisense oligonucleotides or siRNA inhibited cell migration in wounded monolayers. However, specific knockout of AKAP12v1 using CRISPR/Cas9 resulted in increased migration, suggesting that variants 1 and 2 may cooperate in variant-specific, bimodal regulation of cell migration. This notion is supported by the analysis of differential gene expression in endothelial cells lacking AKAP12v1 using RNAseq, which identified more than 4500 differentially expressed genes (DEGs), with Gene Ontology (GO) enrichment analyses and Signaling Pathway Impact Analysis (SPIA) identifying genes linked to cell migration and intercellular junctions. This study is the first to investigate the role of an AKAP12 variant in endothelial cell motility and highlights the complex nature of this signaling scaffold and its regulation of endothelial cell motility.

## Materials and Methods

### Cell Culture

For antisense oligonucleotide and siRNA knockdown experiments, primary human umbilical vein endothelial cells (HUVECs) were obtained from Lifeline Cell Technology (Cat. No. FC0003), and cultured in VascuLife basal medium (Cat. No LL-0002) supplemented with FBS (2%), EnGS (0.2%), rhEGF (0.5 ng/mL), ascorbic acid (50 μg/mL), L-glutamine (10 mM), hydrocortisone hemisuccinate (1μg/mL) and heparin sulfate (0.75 U/mL). The medium was replaced with fresh growth medium three times per week, and the cells were split 1:20 upon reaching 70-90% confluence. Only HUVEC at passages 3-6 were used in this study.

Immortalized HUVECs (ATCC - CRL-4053^™^) were used to create CRISPR/Cas9 mediated AKAP12 knockout cell lines and cultured in VascuLife basal media (Cat. No. LL-0003) supplemented with FBS (2%), basic FGF, VEGF, EGF (5 ng/mL), IGF (15 ng/mL), L-glutamine (10 mM), hydrocortisone hemisuccinate (1 μg/mL), ascorbic acid (50 µg/mL), and heparin sulfate (0.75 U/mL). The cells were passaged 1:5 at subconfluent cell densities and maintained at 37° C in a humidified atmosphere of 5% CO_2_ and 95% air.

### Transfection with knockdown reagents

Antisense and missense oligonucleotides were purchased from Oligos etc. (Wilsonville,OR, USA) as phosphorothioate 20-mers that were either complementary in sequence to a region in the 3’ untranslated region (UTR) of the full-length human AKAP12 transcript (antisense; 5’-CAGTCTCAGCAGCAGCATTC-3’) or a scrambled sequence (missense; 5’-CAGTCTCAGGACCAGCATCT-3’).

Two different siRNAs against AKAP12 (designated siRNA1 and siRNA2) and a non-targeting control or universal negative siRNA (siRNA Control, Cat. No. SIC001) were purchased from Sigma-Aldrich (St. Louis, MO, USA). The sequences of the three siRNAs were as follows: siRNA1, 5’-CGAAACAGCUGUUACCGUA-3’; 5’-UACGGUAACAGCUGUUUCG-3’; siRNA2, 5’-GUAGAAGGUUCCACUGUAA-3’, 5’-UUACAGUGGAACCUUCUAC-3’; and control siRNA, proprietary sequence.

For knockdown experiments using antisense oligonucleotides, HUVEC were incubated in OptiMEM containing Oligofectamine (Invitrogen Life Technologies, NY, Cat. No. 12252-011) and either 0.05μM antisense oligonucleotide, 0.05μM missense oligonucleotide, or no oligonucleotide at 37°C for 4 hr with 5% CO^2^ and then incubated overnight in VascuLife cell culture medium supplemented with VEGF (5ng/ml) and 0.05 μM antisense or missense oligonucleotide. For siRNA experiments, HUVEC were incubated in OptiMEM containing Oligofectamine and 0.25μM siRNA at 37°C for 4 h in a 5% CO_2_ atmosphere, after which the transfection solution was replaced with VascuLife cell culture medium supplemented with VEGF (5ng/mL).

### CRISPR knockout of AKAP12 variant 1

A guide RNA sequence targeting Exon 3 of AKAP12 and containing a protospacer adjacent motif (PAM) (5’-TTAGGGCTCCTTGACCGTTCAGG-3’) was designed, based on the prediction scores for highest ‘on target’ and lowest ‘off target’ events, using an online design tool developed by Dr. Feng Zhang (Massachusetts Institute of Technology). An oligonucleotide with this sequence was then cloned into the plasmid vector pSpCas9(BB)-2A-GFP (PX458) (Addgene) [44] and transfected into subconfluent cultures using Lipofectamine LTX (Invitrogen Cat.15338100) and Plus reagent. Following transfection, GFP-positive cells were isolated by fluorescence-activated cell sorting, cultured until confluent, and then cloned by diluting the cells 1:100 and plating them on 10 cm^2^ dishes to generate single cell-derived colonies. Colonies were selected using trypsin-soaked clonal discs, cultured to 80% confluence, and screened for CRISPR/Cas9 induced mutations.

### Analysis of Genomic DNA for CRISPR/Cas9 Modifications

Clones positive for CRISPR/Cas9 induced indels were identified using a restriction fragment length polymorphism assay (RFLP). Total DNA from the isolated clones was extracted using QuickExtract™ (QE09050), amplified using a NEB Hot Start Taq kit (M0495S) with primers spanning the gRNA target region (5’-CTTGTTTCTGACTTGGTCATGAGGACTG-3’ and 3’-CCAATCCACCACCAGGCAGTATG-5’), digested with HpyCH4III, and run on 2% agarose gels. Clones positive for indels were identified by loss of the HpyCH4III restriction site, and single-base deletions were verified by sequencing amplicons of the RFLP-positive clones. Cell lysates were also analyzed for loss of the AKAP12v1 band on the western blots.

### Effect of cell density and wounding on AKAP12 expression

To determine the effect of cell density on AKAP12 levels, HUVEC were seeded into 25 cm^2^ culture flasks at different cell densities (0.05×10^5^, 0.1 ×10^5^, 0.2×10^5^, 0.6 ×10^5^, 1.2×10^5^ cells/cm^2^) and incubated for 48 h. Following incubation, cells were photographed, harvested, counted, lysed in ice-cold extraction buffer, and analyzed for AKAP12 expression using western blotting, as previously described [34]. Membranes were probed with a rabbit polyclonal anti-AKAP12 antibody (generated previously at the Scripps Research Institute; 1:5000 dilution) [28, 34] and monoclonal alpha actinin antibody (Abcam, MA, Cat No. ab18061; 1:1000 dilution), and the immunoreactive bands were detected using chemiluminescence.

To determine the effect of wounding on AKAP12 levels, monolayers were scratched with a Teflon scraper, labelled for AKAP12 using a monoclonal AKAP12 antibody (2B3-1.1), as described previously [28, 36, 37], and imaged with a Nikon TE300 inverted fluorescence microscope equipped with a Hamamatsu CCD camera. Mean fluorescence intensity was quantified across wounded monolayers on 34 coverslips using ImageJ/Fiji software, with measurements being made at ten 600×50 pixel regions of interest per coverslip.

### Scratch wound migration assays

For antisense and siRNA knockdown studies, HUVECs seeded on tissue culture-treated 8-well plates (Thermo Scientific Nunc, NY) and grown to confluence were wounded by scraping with a 2 mm wide sterile Teflon scraper. Wounded cultures were transfected with antisense oligonucleotides or siRNA as described above and then photographed at 0 h and 18 h post-wounding using a 4X phase contrast objective on a Nikon Diaphot inverted microscope. Reference lines marked on the dishes were used to align the images, and the area between the wound edges at the two time points was measured using ImageJ/Fiji software. Experiments were repeated at least three times.

For experiments using the CRISPR knockout cells, each cell type was seeded in duplicate in 8 well Ibidi µ-slides (Ibidi 80826) (1.5 × 10^5^ cells/per well) and cultured for 3 days to establish a stable monolayer (confirmed by immunolabelling with an anti-VE-cadherin antibody). On day 3, the monolayer was scratched using a silicon scraper, and cultures were then imaged once every 15 min on a Leica DMi8 ThunderImager microscope for 24 h using a stage-top incubation system. The average distance migrated was determined from multiple systematic measurements using the Fiji image analysis software, and the assay was repeated three times.

### Transwell migration assay

For the Transwell migration assay, each cell type was seeded at a density of 3000 cells/well into the top well of a 6.5 mm Transwell chamber containing a PET membrane with 0.8 μm pores (Costar 3464) and cultured under a VEGF gradient created by adding VEGF (35 ng/mL) to the lower chamber. Twenty-four hours later, the cells were scraped from the upper surface of the membranes and the membranes, with the remaining cells, were fixed, stained with Coomassie Blue, and imaged with a Nikon i80 microscope using a systematic random sampling approach [45]. For controls, cultures were not scraped from the Transwell membrane prior to staining. The number of stained cells was quantified using the Fiji image analysis software, and the number of cells that migrated to the lower membrane via invasion was normalized to the total number of cells in the matching control wells.

### RNA Sequencing

RNA was extracted using the RSC Maxwell system with a SimplyRNA kit, and RNA quality was analyzed using a Bioanalyzer system (Agilent Technologies). RNA with an RNA Integrity Number (RIN) greater than seven was used for RNA-Seq library preparation. Ribo-depleted RNA-Seq libraries were prepared using the NEBNext Ultra™ II RNA Library Prep Kit for Illumina (NEB, #E7770). The libraries were sequenced using a HiSeqX instrument, and 40 million 150 bp paired-end reads were generated. Preliminary quality control analysis of FASTQ files was performed using FastQC v0.11.8. The adapters were trimmed using Trimmomatic v0.39 [46]. Reads were aligned to the human genome (hg19) using STAR v2.7.1a [47]. Gene expression was quantified using CuffNorm v2.2.1 [48]. Read counts were summarized using featureCounts v1.4.6 [49]. Differential expression analysis of AKAP12v1 KO vs. WT samples was performed using the R/Bioconductor package DESeq2 v1.24.0 [50]. Genes were considered differentially expressed if they had an FDR ≤ 0.05. Gene Ontology (GO) and Signaling Pathway Impact Analysis (SPIA) enrichment analyses were performed using the R/Bioconductor package clusterProfiler v3.12.0 [51]. Gene Ontology (GO) terms and SPIA pathways were considered enriched if they had a p-value ≤ 0.05, a Benjamini-Hochberg adjusted p-value ≤ 0.05, and a q-value ≤ 0.05.

### Statistical analysis

Statistical analyses were performed using SigmaStat and GraphPad Prism software. Statistical comparisons were performed using Student’s t-test or ANOVA. Comparisons of multiple treatment groups were performed using one-way ANOVA followed by either Tukey’s or Holm-Sidak tests for pairwise comparisons. All statistical tests were two-tailed, with significance reported as follows: * p < 0.05 or p < 0.001. Data are expressed as the mean ±SD.

## Results

### Effect of cell density on AKAP12 expression

Previous studies have shown that AKAP12 expression is low to undetectable in vascular endothelia *<u>in situ</u>*, but is expressed at higher levels in cultured endothelial cells [28, 29]. This suggests that AKAP12 expression in endothelial cells may be regulated by cell density and upregulated under conditions that promote cell migration and proliferation. To investigate this, AKAP12 expression in human umbilical vein endothelial cells (HUVECs) was analyzed using western blotting at different cell densities. As shown in Figure 1A, endogenous AKAP12 expression was lowest at the highest cell density but increased when cell density decreased.

**Figure 1.**
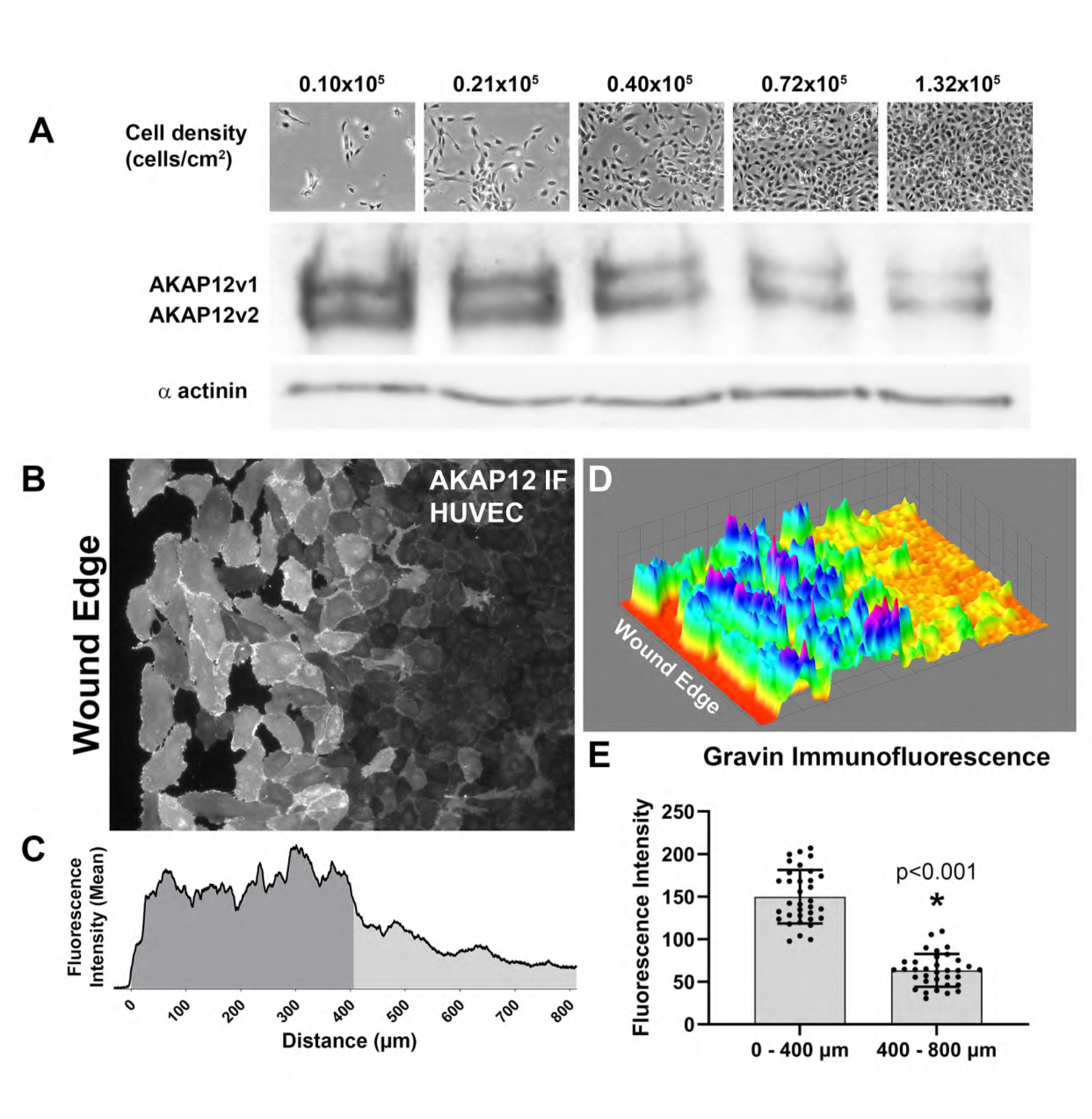
AKAP12 levels are regulated by cell density. A) Phase contrast micrographs and Western blots demonstrating cell density dependent AKAP12 expression in cultured HUVEC. Cells seeded at different densities were harvested and counted 48hr after seeding and analyzed for AKAP12 levels using western blotting. α actinin was used as a loading control. B, C) An representative immunofluorescence micrograph and a corresponding fluorescence intensity plot profile illustrating increased AKAP12 labeling at the wound edge. The micrograph and plot profile are to scale. D) A 3D plot profile of AKAP12 fluorescence levels in the wounded monolayer. D) Bar chart illustrating the mean difference in AKAP12 levels in the dark and light gray regions shown in (C)(N=34; p<0.001).

Based on these results, we examined whether AKAP12 expression was increased in cells at the wound edge in wounded monolayers. Immunofluorescence microscopy of AKAP12 in wounded monolayers showed that AKAP12 was significantly upregulated in a layer of cells approximately 400 µm wide at the wound edge (Fig. 1 B, C, D, E). This change in expression was also accompanied by a change in AKAP12 distribution, with AKAP12 being concentrated at junctions in confluent regions of the monolayer but distributed to regions reminiscent of lamellipodia and membrane ruffles in cells at the edge of the wound.

### Effect of AKAP12 knockdown on wound closure

To test the hypothesis that AKAP12 plays a role in endothelial cell wound healing, the effect of AKAP12 knockdown on wound closure in cultured HUVEC was determined using an antisense phosphorothioate oligonucleotide targeting the 3’ UTR of the message (exon 6) and two siRNAs targeting different regions in the coding sequence of the message (exon 5). Both the antisense oligonucleotide and siRNA treatments significantly reduced AKAP12v1 and AKAP12v2 levels (Fig. 2A), and treatment with either the antisense oligonucleotide or siRNA significantly reduced the distance that cells at the wound edge migrated following a scratch wound (Fig. 2B, C). Surprisingly, while siRNA#1 had the greatest knockdown efficiency, its effect on cell migration was less than that of siRNA#2 or antisense oligonucleotide. This result was repeated through multiple trials and suggests that endothelial cells may be sensitive to fine-tuning of AKAP12 expression during cell motility.

**Figure 2.**
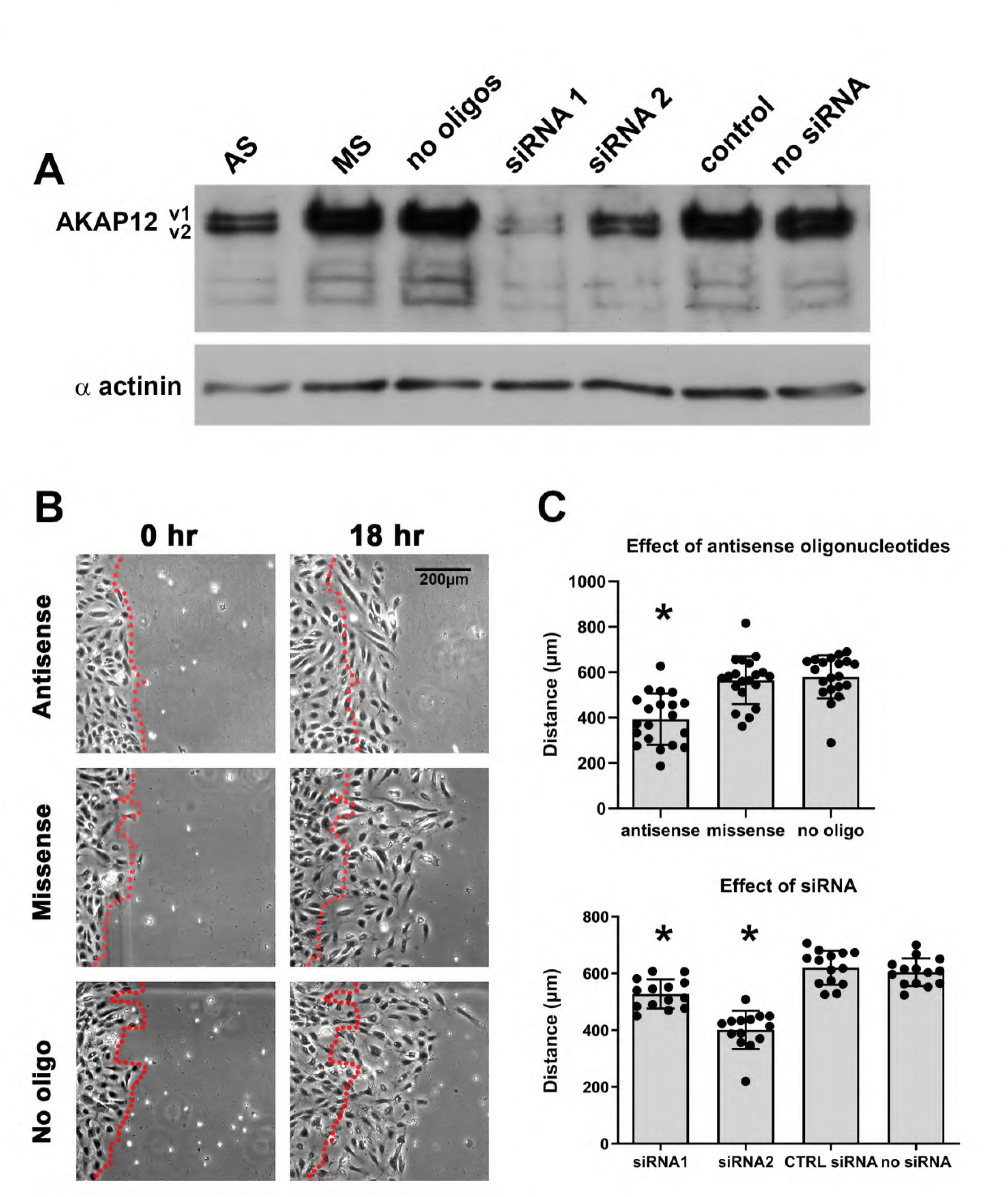
Effect of AKAP12 knockdown on cell migration. A) A western blot illustrating the reduction in AKAP12 levels after either antisense oligonucleotide or siRNA treatment. B) Phase contrast micrographs of wounded monolayers showing a decrease in migration by HUVECs treated with an antisense oligonucleotide. C) Bar graphs illustrating the decrease in HUVEC migration after treatment with either the antisense oligonucleotide or the siRNAs (N=3; p<0.05).

### Endothelial cells expressing only AKAP12 variant 2 showed increased invasion and migration

Given that knockdown of both AKAP12 variants with antisense oligonucleotides and siRNA inhibited endothelial cell migration, we wanted to determine whether the elimination of one of the variants, AKAP12v1, would similarly affect HUVEC migration. Previous studies on the role of AKAP12 in migration have primarily examined the effects of overexpressing the variant 1 coding sequence in tumor cells using plasmid vectors [52–55]. Only one study has reported the role of AKAP12v2 in cell migration [43]. Here, we used CRISPR/Cas9 to generate HUVEC cell lines lacking AKAP12v1 while maintaining the expression of variant 2. This was accomplished by transfecting HUVEC-TERT2 cells with an SpCas9 plasmid containing a gRNA targeting exon 3 and screening for clones lacking an HpyCH4III site near the PAM site, as described in the Methods and Materials section (Fig. 3A, B). An RFLP assay using genomic DNA identified three clones that were positive for the loss of the HpyCH4III restriction enzyme site spanning the predicted location of indels arising from NHEJ repair of a CRISPR-Cas9 dependent double stranded break (Fig. 3D). These clones were designated AKAP12v1 KO Clone1, Clone2, and Clone3. Sequencing of PCR products spanning this region showed a homologous single nucleotide deletion of a C base, four bases from the PAM site, thereby resulting in a frameshift mutation and the introduction of a predicted stop codon five codons downstream of the break site (Fig. 3B, C). RNA-Seq analysis revealed that there was a significant decrease in the amount of AKAP12v1 transcripts in the KO clones compared to the WT control, while there was no significant difference in the AKAP12v2 transcripts between WT and KO clones (Fig. 6B). Western blotting of whole-cell lysates confirmed the loss of AKAP12v1 in the modified clones compared to that in the WT control cells (Fig. 3E, F). Taken together, we successfully created three AKAP12v1 KO cell lines, all of which had a single nucleotide deletion that resulted in a premature stop codon, a significant decrease in AKAP12v1 transcript, and a complete loss of the AKAP12v1 protein, possibly via nonsense-mediated decay of the aberrant v1 transcript [56]. An effort to create AKAP12v2 KO mice was also undertaken using the same approach. However, despite several attempts, we were unsuccessful in creating this knockout because the coding region unique to AKAP12v2 was very short and lacked regions that could be targeted to produce single indels that would create a nonsense sequence or premature stop codon.

**Figure 3.**
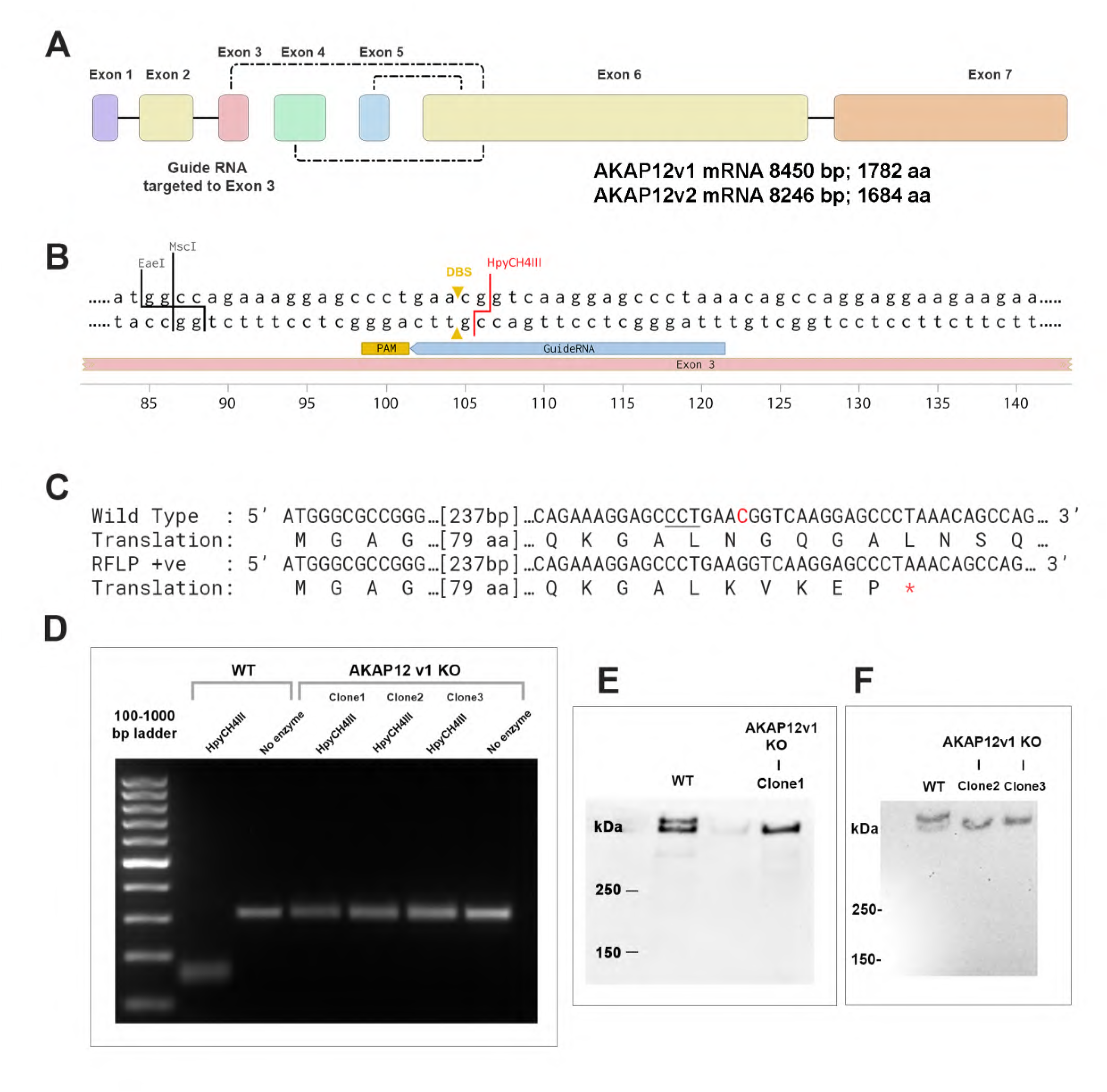
Creation of an AKAP12v1 knockout using CRISPR-Cas9 with non-homologous recombination. A) A diagram illustrating the exon organization of human AKAP12 and the exon combinations that give rise to the three AKAP12 variants. As illustrated here, exons 1, 2, and 3 combine with exons 6 and 7 to give rise to AKAP12v1, exon 4 combines with exons 6 and 7 to give rise to AKAP12v2 and exon 5 combines with exons 6 and 7 to give rise to AKAP12v3. AKAP12v3 is not expressed in vascular endothelial cells. B) A diagram illustrating the region in exon 3 targeted by the guide RNA used to produce the AKAP12v1 knockout. The HpyCHIII site used for the RFLP analysis and the location of the double stranded break (DBS) generated by CRISPR-Cas9 are shown here. C) A diagram illustrating both the wild type and CRISPR-Cas9 modified (RFLP +ve) AKAP12v1 mRNA coding sequences showing the location of the PAM site (<u>CCT</u>), the location of the single base deletion (marked in red), and the introduction of a stop codon in the modified AKAP12v1 sequence. D) An agarose gel illustrating the results of an RFLP analysis and identification of three knockout cell lines lacking the HpyCHIII restriction site. E, F) Western blots illustrating loss of the larger AKAP12v1 band in the knockout cells.

The effect of AKAP12v1 knockout on endothelial cell migration was assessed using both transwell migration and scratch wound assays. Surprisingly, KO of AKAP12v1 caused substantial elevation in transwell migration within 24 h compared to WT control cells (Fig. 4A,B). Similarly, AKAP12v1 knockout cells migrated further than wild-type cells in a scratch wound assay. Analysis of three independent scratch wound experiments showed that all three AKAP12v1 KO clones migrated significantly further than WT cells at 24 h, albeit with some variability (Fig. 5A, C; Supplementary Movie 1). The movement of individual cells at the edge of the monolayer was tracked using a manual tracker and chemotaxis plugin, and all three AKAP12v1 KO clones showed significantly higher single-cell velocity and single-cell accumulated distance values compared to WT cells (Fig. 5B, C).

**Figure 4.**
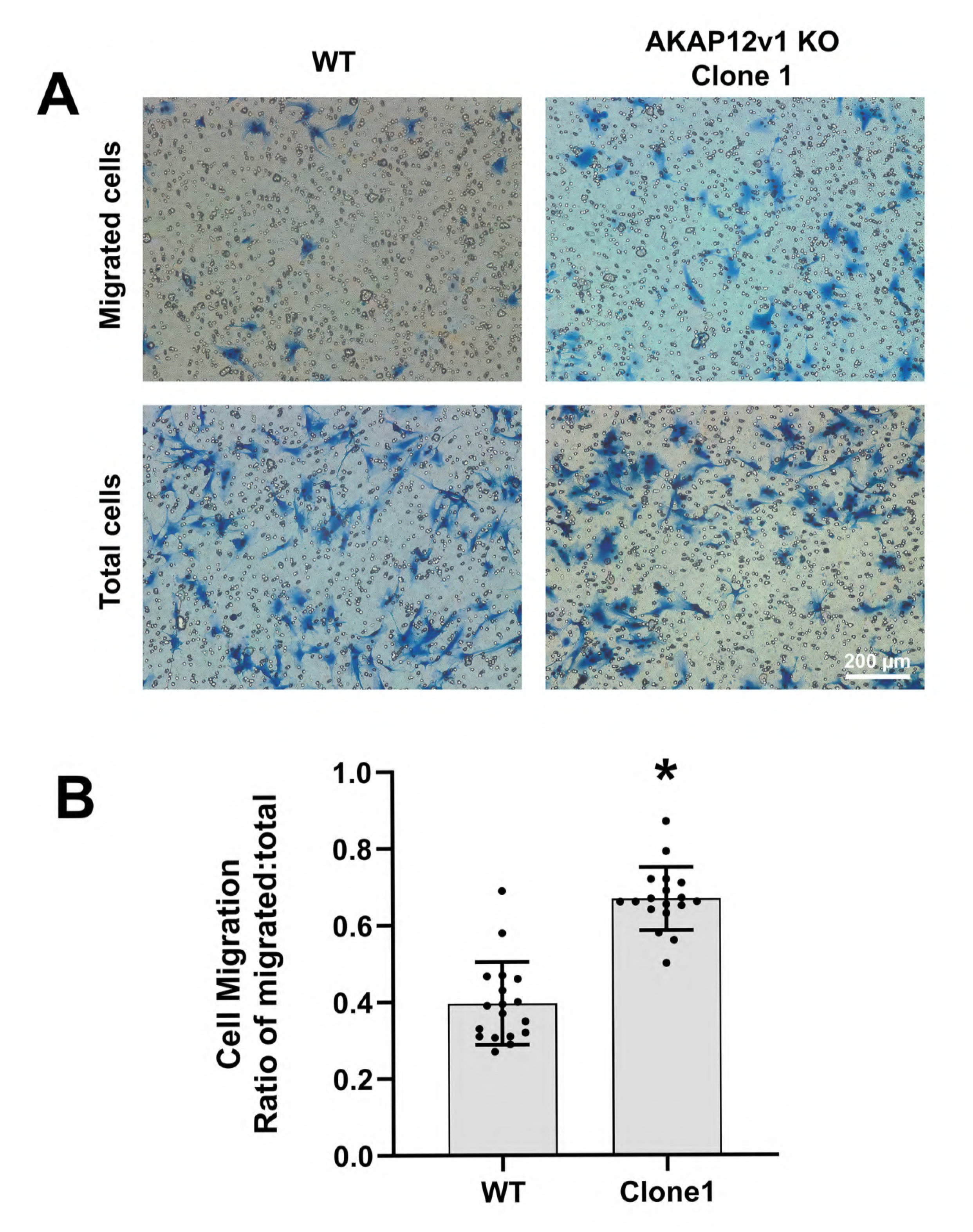
A Transwell migration assay showing increased migration by one of the AKAP12v1 knockout cell lines (Clone1) compared to wild type (WT) cells. A) Micrographs of toluidine blue stained cells showing the number of knockout and WT cells on the underside of the Transwell membrane after 24 hours incubation. B) Quantification of WT and knockout cell migration expressed as the ratio of cells that have passed through the membrane to the total number of cells at the beginning of the assay. N=3; p<0.05.

**Figure 5.**
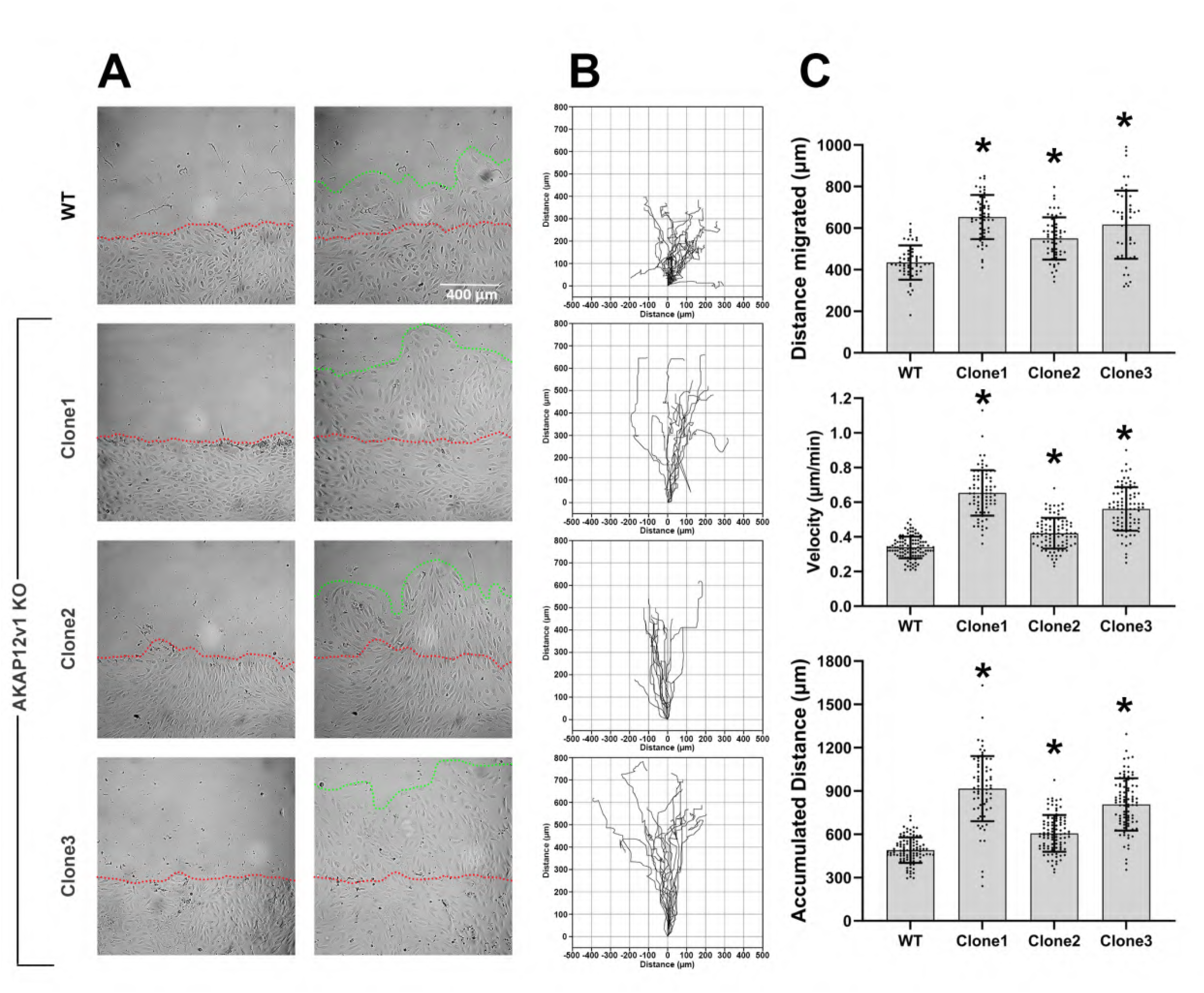
Scratch wound migration assay comparing the three knockout cell lines to WT cells. A) Phase contrast images illustrating migration distances of each of the cell lines. B) Tracings of a sample of individual cells at the wound edge illustrating their migration pattern. C) Graphs illustrating the mean overall migration distance for each cell line (top), the mean velocity of the individual cells tracked in B (middle), and the accumulated migration distance the individual cells tracked in B (bottom). N=3; p<0.05.

### RNA-seq analysis of AKAP variant 1 knock out cells reveal signaling pathways related to migration

Having observed that endothelial cells lacking AKAP12v1 but expressing AKAP12v2 migrated significantly faster than WT cells, we postulated that this would be reflected in their transcriptomic landscape. To test this, we performed bulk RNA sequencing of all three KO clones using confluent cells in culture and compared their gene expression profiles to that of WT cells. As demonstrated in the principal component analysis (PCA) and Venn diagram (Fig. 6A, C), the transcriptome of AKAP12v1 KO clones differed from that of WT cells. Although the transcriptomes of the three clones showed some variability, ∼70% of the total differentially expressed genes (DEGs) were common between any two clones and 40% were common to all three clones (Fig. 6C). Despite using gRNA with low off-target predictions, this clonal variability could be due to the intrinsic heterogeneity of the WT cells [57]. As a result, only a subset of 40% common genes was used in Gene Ontology (GO) enrichment analysis and Signaling Pathway Impact Analysis (SPIA) to better understand the changes in the transcriptomic landscape when AKAP12v1 was knocked out.

**Figure 6.**
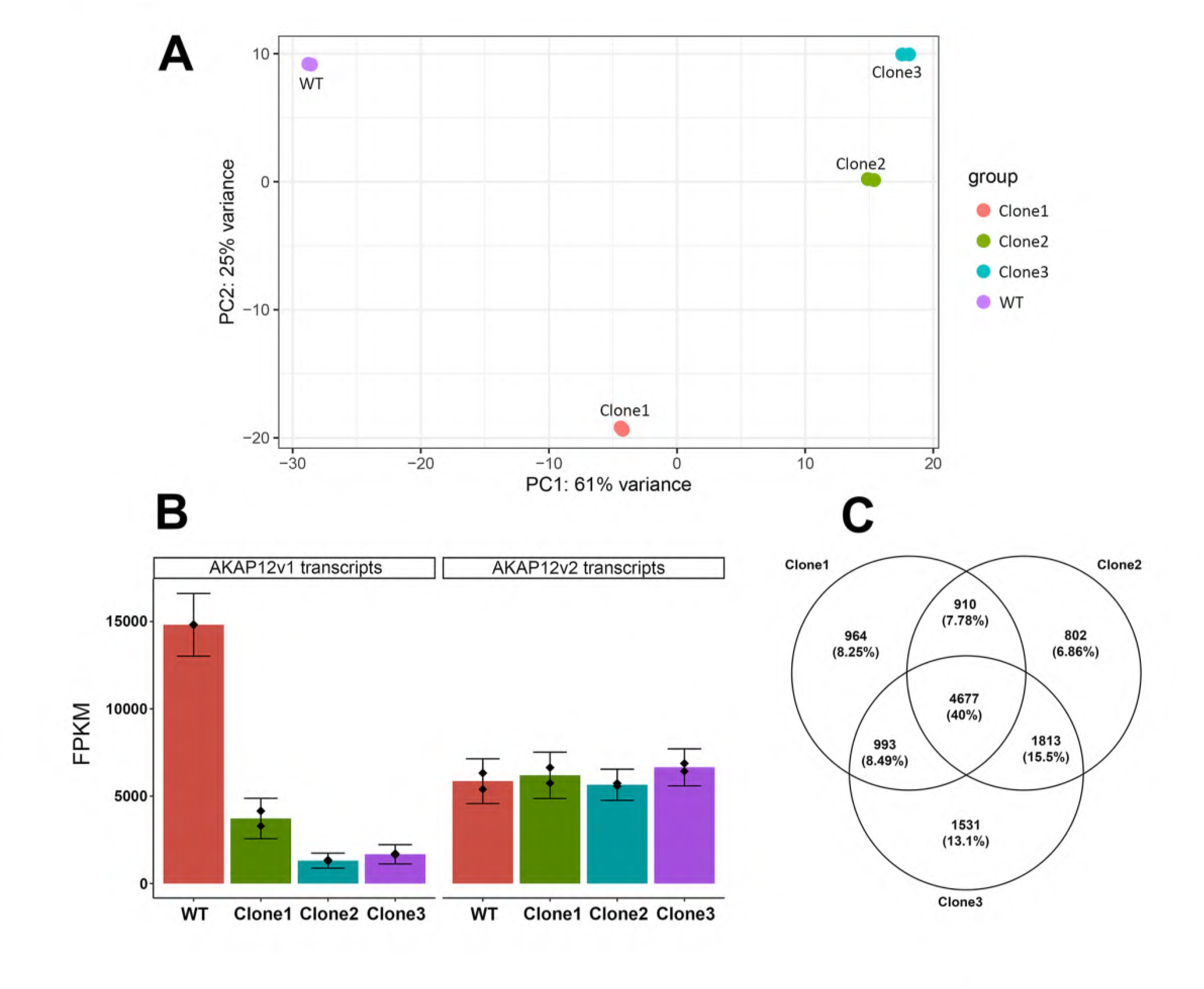
Comparison of AKAP12v1 KO and WT HUVEC transcriptomes. A) PCA plot illustrating that all three AKAP12 v1 KO clones are significantly different from WT cells, but with some variability among Clones 1, 2, and 3. Results are from two sets of samples. B) FPKM plot of AKAP12v1 and v2 transcripts from the RNA-seq data showing reduced AKAP12v1 transcripts in the three KO clones compared to WT cells, but no difference in the relative number of AKAP12v2 transcripts between the WT and KO cell clones. Results are from two sets of samples. Bars = confidence intervals. C) Venn diagram showing the number of unique and common DEGs associated with the three KO clones. 40% of the DEGs are common to all three AKAP12 v1 KO clones.

Consistent with the western blot results described above, all three clones of the AKAP12v1 KO showed decreased AKAP12 transcript levels (Fig. 6B), likely due to the presence of CRISPR-mediated indels in the AKAP12v1 mRNA inducing nonsense-mediated decay of the transcripts [57]. In contrast, there was no significant difference in AKAP12v2 transcript levels between WT and KO clones (Fig. 6B). In addition, we queried the dataset for the differential expression of known signaling molecules that are part of the AKAP12 signalosome; however, no significant changes were observed.

Endothelial cell migration is well characterized in the literature and is regulated by a network of interconnected signaling pathways that are coupled to molecular functions (MF), cellular components (CC), and biological processes (BP) in the cell. As expected, Gene Ontology analysis of the transcriptome of AKAP12v1 KO clones revealed significant MF, CC, and BP changes known to be involved in endothelial cell migration (Supplementary Tables 1, 2, and 3). Highly significant BP elements include the regulation of actin filament-based processes, regulation of actin cytoskeleton organization and assembly, cell-substrate adhesion, focal adhesion assembly, and signaling pathways related to these processes. Similarly, the most significant CC and MF elements were related to cell-substrate interactions and characteristics associated with cell motility (e.g., leading edge, lamellipodia, and cell cortex).

SPIA enrichment analysis, which combines the evidence obtained from classical enrichment analysis with actual perturbation of a given pathway under a given condition, showed that the DEGs common to all three AKAP12v1 KO clones were assigned to more than 100 pathways that may have been modified. Of these pathways, 17 had Bonferroni-adjusted global p-values <0.05, with 6 pathways predicted to be activated and 11 predicted to be inhibited (see additional data under Supplementary Data). Among these pathways, several were closely related to endothelial cell migration; VEGF signaling was predicted to be activated, while focal adhesion, extracellular matrix (ECM)-receptor interaction, regulation of actin cytoskeleton, MAPK signaling, and tight junction pathways were predicted to be inhibited (Supplementary Table 4). Further analysis of these six predicted pathways revealed that only a small number of DEGs associated with each pathway were significantly upregulated by >1. 5 fold while most genes were significantly downregulated (<-1.5)(Fig. 7A). As anticipated, the analysis of these pathways for overlapping DEGs revealed that most were associated with ECM interactions, focal adhesions, and actin cytoskeletal regulation pathways (Fig.7B). The majority of these DEGs, which included several members of the integrin family, several types of collagen, fibronectin, vitronectin, myosin light chain 9, myosin light chain kinases, guanine nucleotide exchange factor *ARHGEF6* (*α*-PIX), and TGF*β*2, were significantly downregulated, while some DEGs, including *EPB41*, *vWF*, *LAMA2*, *CD36*, *PIK3CG* and *NCKAP1L* were upregulated to varying degrees. These resulting transcriptomic changes are consistent with the effect of AKAP12v1 loss on cell migration in transwell invasion and wounding experiments.

**Figure 7.**
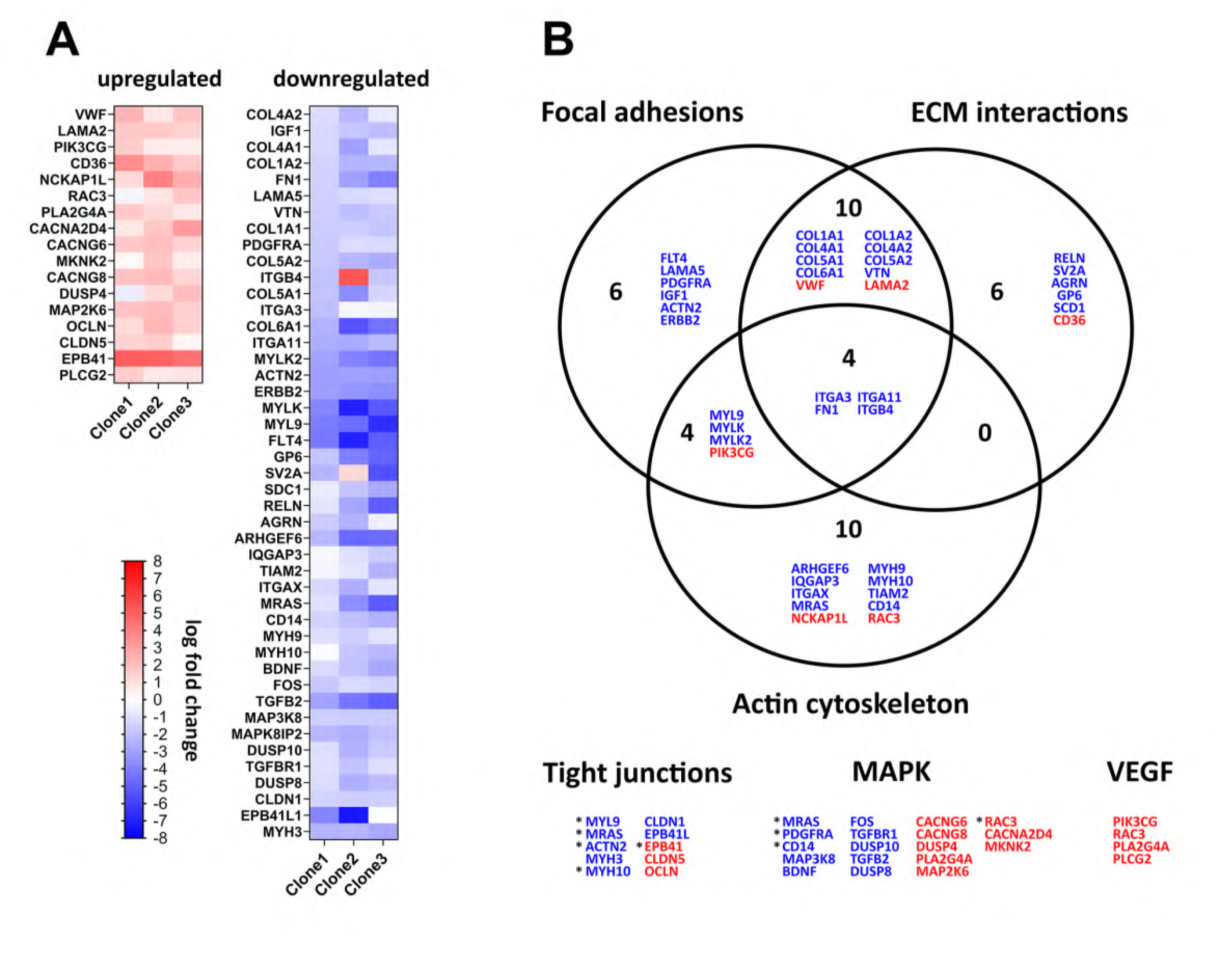
Diagrams highlighting the DEGs (upregulated or downregulated) common to the KO clones that were linked by Gene Ontology and SPIA enrichment analysis to 6 pathways closely related to endothelial cell motility. A) Heat map charts illustrating the upregulated (red) and downregulated (blue) DEGs related to focal adhesions, ECM interactions, actin cytoskeletal regulation, tight junctions, MAPK, and VEGF pathways. The log fold change in expression is indicated by the scale bar. B) Venn diagram showing that many of the DEGs common to focal adhesions, ECM interactions, and actin cytoskeletal regulation were shared among these three pathways. DEGs associated with the tight junctions, MAPK, and VEGF pathways had little overlap with each other or the other three pathways. Upregulated genes are marked in red; downregulated genes are marked in blue. Asterisks beside the genes linked to tight junctions, MAPK, and VEGF pathways identify those that are shared with another pathway.

## Discussion

In this study, we report that the expression of AKAP12v1 and v2 in endothelial cells is dependent on cell density, with expression being low in confluent endothelial cell monolayers but upregulated in subconfluent endothelial cell cultures and in endothelial cells at the wound edge in wounded monolayers. Knockdown of both variants with either an antisense oligonucleotide or siRNAs inhibited endothelial cell migration, but the effect of knockdown appeared to be biphasic, with higher levels of knockdown having less effect on migration than moderate levels of AKAP12 reduction. Following these experiments, we investigated the effect of deleting AKAP12v1 using CRISPR/Cas9 and found that endothelial cells expressing only AKAP12v2 showed higher rates of migration than wild-type HUVEC, suggesting that the two variants may have different roles in regulating the signaling pathways associated with cell motility.

The demonstration that AKAP12 expression in endothelial cells is cell density-dependent and upregulated at the wound edge during wound repair extends our previous work [29] and confirms the results of a recent study by Benz et al. [58], who also reported elevated AKAP12 expression in endothelial cells at the wound edge. A number of transcriptomic studies have shown similar effects of wounding on AKAP12 expression in endothelial cells and other epithelia [59, 60]. For instance, supplementary transcriptomic data linked to a study investigating endothelial cell proliferation during wound repair of the aorta in mice showed upregulation of several genes, including AKAP12 [59], and a transcriptomic study by Fitsialos et al. [60] reported AKAP12 among the genes upregulated in response to wounding in a keratinocyte wound repair model. The regulatory pathways responsible for AKAP12 upregulation in vascular endothelial cells are still poorly understood, but several studies have identified several potential players. These include HIF1α [41], VEGF [61], HGF [61], HDAC7 [62], angiopoetin-1 [63], and VE-cadherin [64]. In particular, changes in VE-cadherin expression may be an initiating signal leading to the upregulation of AKAP12 in response to the wounding of endothelial cell monolayers. Morini et al. [64] showed that transfection of VE-cadherin into VE-cadherin null endothelial cells resulted in the upregulation of several genes linked to junction stabilization but also led to the downregulation of several genes, including AKAP12. This effect of VE-cadherin expression on AKAP12 is consistent with the cell density-dependent changes in AKAP12 protein levels reported here. VE-cadherin is well known to be elevated and localized at cell junctions in confluent monolayers, where we observed low AKAP12 expression, but is reduced and lost at cell margins in dispersed low-density cultures and at the edge of wounded monolayers, where we observed AKAP12 expression to be elevated.

Cell density-dependent changes in AKAP12 expression were also accompanied by changes in AKAP12 distribution. AKAP12 is concentrated at cell margins in confluent monolayers and has been linked to the regulation of junctional permeability [41, 42]. However, AKAP12 expression in cells at the wound edge was elevated over the entire cell surface and concentrated at the ruffles and lamellipodia, regions associated with endothelial cell motility, a finding also noted by Benz et al. [58]. These changes in AKAP12 expression and distribution suggest that this signaling protein scaffold may play a dynamic regulatory role in endothelial cell activity by regulating cellular junctions in confluent endothelial cell monolayers through localization of signaling proteins such as PKA to cell margins [41, 42], but plays a role in cell motility at the wound edge and in dispersed cells by localizing signaling proteins (e.g. PKA, PKC, and Src) to lamellipodia, where they may control actin dynamics and cell-substrate interactions [24, 29, 31, 32, 34, 36, 37, 58].

Consistent with the finding that AKAP12 expression increases at the wound edge in wounded monolayers, AKAP12 knockdown with antisense oligonucleotides and siRNAs inhibited wound healing in endothelial cell monolayers, confirming the results of a previous study by Benz et al. [58], who used endothelial cells from AKAP12 knockout mice to investigate the role of AKAP12 in endothelial cell migration. Interestingly, the effect of knockdown appeared to be biphasic, with partial knockdown of AKAP12 resulting in greater inhibition of migration than the almost complete AKAP12 knockdown seen with one of the siRNA treatments (siRNA1). This result was consistent in multiple trials suggesting that endothelial cells may be sensitive to fine-tuning of AKAP12 expression during cell motility. Benz et al. [58] reported that complete knockout of AKAP12 in mouse endothelial cells inhibited migration, but this was observed in the presence of VEGF at concentrations higher than those used in the current study. However, these results clearly show that AKAP12 plays an important role in due to the fact that 1) AKAP12 interacts with multiple signaling proteins known to play a role in cell motility, 2) cell motility is controlled by multiple input signals, and 3) two AKAP12 variants are expressed in endothelial cells simultaneously.

To address this, we examined the role of AKAP12v2 alone in endothelial cell migration. Previous AKAP12 research related to cell motility has focused almost entirely on AKAP12v1, with only limited consideration of AKAP12v2 [43]. Endothelial cells expressing only variant 2 were generated by knocking out AKAP12v1 using the CRISPR-Cas9 technology. Western blotting confirmed the loss of AKAP12v1 in the knockout cells, and RNA-seq data confirmed a significant reduction in AKAP12v1 transcripts. Loss of AKAP12v1 resulted in increased invasion in a Transwell membrane assay and increased cell migration in a scratch wound assay. Consistent with this, RNA-seq analysis identified transcriptomic changes in AKAP12 v1 KO clones that predicted changes in pathways such as focal adhesions, tight junctions, cell substrate adhesion, and regulation of the actin cytoskeleton, which reflected cellular events associated with endothelial cell migration, barrier function, and angiogenesis. This effect of knocking out AKAP12v1 expression on endothelial cell migration is consistent with a previous study by Finger et al. [43], in which they linked elevation of AKAP12v2 levels in melanoma cells in response to hypoxia to elevated motility. Based on these results, we propose that AKAP12 variants each have an individual role but work together in a balanced manner to regulate the complex and interconnected signaling networks controlling endothelial cell motility and migration.

Structural differences between the two AKAP12 variants support the hypothesis that they have individual roles in regulating endothelial cell motility. For instance, both variants possess several membrane localization motifs that direct them to the plasma membrane [24, 34, 36–38], but variant 1 possesses a myristoylation motif at the N-terminus that is absent in variant 2. Previous work has shown that the membrane localization and trafficking properties of variant 1 are altered when the myristoylation domain is deleted [34]; thus, variant 1 may localize to a different membrane microdomain or have different membrane-binding characteristics than variant 2. If so, this would likely mean that the targeting of signaling proteins bound to these variants would also differ. Both variants also possess an Src-binding domain that has been linked to Src-dependent FAK regulation [32], but variant 1 possesses an additional Src-binding domain that has been linked to the regulation of β2-adrenergic receptor internalization and recycling [31]. While it is not clear how these differences translate into different roles for each variant in cell adhesion and motility, extension of a model proposed by Su et al. [32] could account for the finding that cell migration is elevated in cells lacking variant 1. In their model, AKAP12 sequesters a pool of Src/Cav-1 in lipid rafts and regulates normal focal adhesion turnover by maintaining a dynamic balance between this pool and Src/Cav-1 at focal adhesion sites. Loss of AKAP12 shifts this balance toward localization of Src/Cav-1 at focal adhesions, resulting in increased Src-dependent activation of FAK, increased focal adhesion turnover, and increased migration/invasion. Given that AKAP12 exists as two variants with potentially different membrane localization characteristics, it is reasonable to reenvision this model such that variant 1 sequesters Src in the lipid raft pool, and variant 2 plays a role in scaffolding Src at focal adhesions. Deletion of variant 1 would disrupt the balance between the two Src pools, leading to increased focal adhesion turnover and increased motility. Such a model could also account for other AKAP12 dependent signaling events that control focal adhesion and lamellipodia dynamics, such as PKA-dependent phosphorylation of Rho [65] or VASP [41, 58]. By scaffolding PKA at lipid rafts and non-raft membrane regions, variants 1 and 2 may establish separate AKAP12-PKA pools in dynamic balance. Deletion of variant 1 would alter this balance, which in turn could lead to elevated phosphorylation of PKA targets, resulting in increased lamellipodial dynamics and cell motility. Further studies to demonstrate the differential membrane localization of the two AKAP12 variants and changes in signaling protein binding to each of the variants in response to elevated cellular motility will be critical to verify this model.

Transcriptomic profiling via bulk RNA sequencing confirmed that AKAP12v1 was significantly downregulated and that the resulting transcriptomic changes were consistent with the results of endothelial cell migration and invasion experiments. ECM interactions, focal adhesions, and actin cytoskeleton pathways known to regulate migration were most affected in knockout clones. Of the genes involved in these pathways, a higher proportion were downregulated and included genes for several integrin subunits, collagen subtypes, *LAMA5*, vitronectin, and fibronectin 1, all of which have well-known roles in cell adhesion and motility [66–68]. In contrast, the upregulated genes linked to these pathways included *vWF*, *LAMA2*, *CD36*, *PIK3CG* and *NCKAP1L*. *LAMA2* has been shown to play a role during development and in processes such as cell adhesion, migration, and blood brain barrier integrity [69, 70]. *NCKAP1L* is part of the WAVE complex, which is a critical regulator of the actin cytoskeleton and lamellipodia formation during migration [71, 72]. vWF, a well-studied protein in endothelial cells, promotes transendothelial cell migration [73] and regulates endothelial cell proliferation, migration, and angiogenesis via VEGFR2 signaling [74]. CD36 is a lipid scavenger receptor that has pro-metastatic functions in several cancers [75] and has been shown to increase the migration of endothelial cells in an AMPK-dependent manner [76]. *PIK3CG*, which encodes the PI3Kγ (phosphatidylinositol 3-kinase gamma) protein, plays a significant role in regulating endothelial cell migration, particularly during angiogenesis, by activating signaling pathways that promote migration and adhesion to the extracellular matrix [77, 78]. Interestingly, EPB41, which was associated with the tight junction pathway in this study, showed the highest level of upregulation in variant 1 KO cells. This protein, also called protein 4.1R, is well known to link the actin cytoskeleton to the membrane and membrane-associated proteins, and has been shown in a limited number of studies to play a role in epithelial cell migration [79, 80]. Taken together, these transcriptomic changes, which indicate changes in pathways linked to cell motility, are consistent with the elevated motility observed in AKAP12v1 KO clones in this study. A more detailed analysis of the DEGs and pathways identified in this study will be necessary to reach a more comprehensive understanding of how AKAP12 variants regulate endothelial cell motility.

The past 20 years have witnessed significant advances in our understanding of the complexities and subtleties underlying the regulation of cell motility. However, this study is the first to provide evidence that AKAP12 variants may play different roles in endothelial cell motility and wound repair. In this study, we introduced an investigative approach wherein CRISPR-Cas9 genome editing was used to identify the functional significance of AKAP12v1 loss in endothelial cells. The results of this study suggest that AKAP12v2 and AKAP12v1, which are regulated by different promoters, may play different roles but function in concert to regulate endothelial cell migration. However, further studies will be required to determine the extent to which AKAP12 variant expression, cellular distribution, and interactions with their signaling protein partners control endothelial cell motility. In turn, this knowledge may lead to an understanding of how manipulation of the expression and activity of the two variants can be used to control vascular repair processes.

## Supporting information

Supplemental Additional RNAseq data

Supplemental original WB

## Acknowledgement

This work was supported in part by pilot grants funded by NIH P30GM103329 and NIH P30GM103329, and funds from NIH P20GM113123. In addition, the authors acknowledge Sarah Abrahamson (Senior Research Specialist - UND Imaging Core), Steve Adkins ( Research Specialist – UND Flow Cytometry Core), and Danielle Perley and Brett McGregor (Research specialists – UND Genomics Core) for technical assistance with the study. Moreover, the authors acknowledge the use of the UNDSMHS Imaging and Image Analysis Core Facility supported by NIH grants P20GM113123 and U54GM128729, the North Dakota Flow Cytometry and Cell Sorting Core supported by NIH grant P20GM113123, and the Genomics Core funded in part by NIH grants U54GM128729 and P20GM104360-06A1.

## Statement of Ethics

Study approval statement: The study protocol was reviewed and approved by the University of North Dakota Institutional Biosafety Committee (approval number [IBC-201801-009]).

Consent to participate statement: This study did not use human participants and therefore did not require informed consent.

## Author Contributions

The authors confirm their contribution to the paper as follows: study conception and design: Ashrifa Ali, Bhaskar Roy, Micah Schott, Bryon Grove; data collection: Ashrifa Ali, Bhasker Roy, Micah Schott ; analysis and interpretation of results: Asrhifa Ali, Bhaskar Roy, Micah Schott, Bryon Grove; draft manuscript preparation: Ashrifa Ali, Bryon Grove. All authors have reviewed the results and approved the final version of the manuscript.

## Data Availability Statement

All data generated or analyzed during this study are included in this article and its supplementary material files. Further inquiries can be directed to the corresponding authors.

The RNA-seq data discussed in this publication have been deposited in NCBI’s Gene Expression Omnibus (Edgar et al., 2002) and are accessible through GEO Series accession number GSE285043 (https://www.ncbi.nlm.nih.gov/geo/query/acc.cgi?acc=GSE285043).

## Supplementary data

Supplementary movie 1. A movie illustrating the difference in migration rate between wounded WT and AKAP12v1 KO HUVEC. 15 min/frame for 24 hours.

Supplementary Table 1. A list of biological processes that may be perturbed in endothelial cells as a result of knocking out AKAP12v1. The majority are related to the regulation of cytoskeleton and cell-substrate adhesion. fDR = false discovery rate.

Supplementary Table 2. Cellular characteristics that may be perturbed in endothelial cells as a result of knocking out AKAP12v1. Consistent with the increase in cell migration observed in the scratch wound assay, the cellular characteristics predicted the perturbation of focal adhesions, cell substrate junctions, and actin cytoskeletal changes. fDR = false discovery rate.

Supplementary Table 3. A list of the molecular functions associated with perturbation by knocking out AKAP12v1 in endothelial cells. These functions include actin binding, cell-cell interactions, cell-substrate interactions, and signaling pathways, all of which are known to play a role in cell motility. fDR = false discovery rate.

Supplementary Table 4. Signaling Pathway Impact Analysis (SPIA). A list of perturbed pathways resulting from knocking out AKAP12v1 in endothelial cells. Consistent with the outcome of the phenotypic assays, the focal adhesion and actin cytoskeleton regulation pathways were predicted to be significantly perturbed by more than 70 DEGs participating in this pathway. ECM-receptor interactions and tight junctions were also predicted to be perturbed in AKAP12v1 KO cells. pSize = number of genes in the pathway; NDE = number of differentially expressed genes; pNDE = hypergeometric probability of observing NDE differentially expressed genes in the pathway by chance; tA = observed value of the perturbation score; pPERT = bootstrap probability associated with tA; pGFDR = adjusted pG using the false discovery rate correction; pGFWER = adjusted pG using family-wise error rate (Bonferroni); Status = inhibition/activation according to the -/+ sign of tA.

Additional supplementary data. Complete dataset for biological process analysis, cellular component analysis, molecular function analysis, SPIA enrichment, and differentially expressed genes common to the three AKAP12v1 KO clones. Pathways highlighted in green under the SPIA enrichment tab correspond to 17 pathways with Bonferroni-adjusted global p-values (pGFWER) <0.05.

Fig. 1 original western blot. Cropped regions marked in red.

Fig. 2 original western blot. Croppepd regions marked in red.

Fig 3E original blot with molecular weight markers showing 250 kDa and 150 kDa locations on the western blot). Cropped region marked in red. Original blot for Fig.3F is not available.

