## Supplemental tables 1-4 for "AKAP12 variant 1 knockout enhances vascular endothelial cell motility"

Supplementary Table 1. Gene Ontology: Biological Processes

| ID | Description | GeneRatio | fDR |
|---|---|---|---|
| GO:0032970 | regulation of actin filament-based process | 167/4018 | 4.07E-18 |
| GO:0032956 | regulation of actin cytoskeleton organization | 152/4018 | 4.72E-18 |
| GO:0043087 | regulation of GTPase activity | 181/4018 | 1.48E-13 |
| GO:0031589 | cell-substrate adhesion | 142/4018 | 8.94E-13 |
| GO:0043547 | positive regulation of GTPase activity | 151/4018 | 1.02E-10 |
| GO:0032271 | regulation of protein polymerization | 94/4018 | 2.49E-10 |
| GO:0051017 | actin filament bundle assembly | 69/4018 | 1.30E-08 |
| GO:0008064 | regulation of actin polymerization or depolymerization | 77/4018 | 1.58E-08 |
| GO:0010810 | regulation of cell-substrate adhesion | 88/4018 | 1.67E-08 |
| GO:0045785 | positive regulation of cell adhesion | 141/4018 | 3.79E-08 |
| GO:0061572 | actin filament bundle organization | 69/4018 | 3.79E-08 |
| GO:0007044 | cell-substrate junction assembly | 49/4018 | 4.19E-08 |
| GO:0150115 | cell-substrate junction organization | 49/4018 | 4.19E-08 |
| GO:0001666 | response to hypoxia | 125/4018 | 3.67E-07 |
| GO:0034446 | substrate adhesion-dependent cell spreading | 47/4018 | 1.16E-06 |
| GO:0043149 | stress fiber assembly | 46/4018 | 1.64E-06 |
| GO:0048041 | focal adhesion assembly | 40/4018 | 2.23E-06 |
| GO:0007266 | Rho protein signal transduction | 78/4018 | 2.31E-06 |
| GO:0034329 | cell junction assembly | 134/4018 | 4.84E-06 |
| GO:0061028 | establishment of endothelial barrier | 24/4018 | 1.53E-05 |
| GO:0051403 | stress-activated MAPK cascade | 97/4018 | 3.32E-05 |
| GO:1901888 | regulation of cell junction assembly | 70/4018 | 5.25E-05 |
| GO:0046578 | regulation of Ras protein signal transduction | 82/4018 | 8.74E-05 |
| GO:0006469 | negative regulation of protein kinase activity | 81/4018 | 9.54E-05 |
| GO:0045216 | cell-cell junction organization | 67/4018 | 9.81E-05 |

Supplementary Table 2. Gene Ontology: Cellular Components

| ID | Description | GeneRatio | fDR |
|---|---|---|---|
| GO:0005925 | focal adhesion | 210/4138 | 9.75E-41 |
| GO:0030055 | cell-substrate junction | 211/4138 | 3.79E-40 |
| GO:0031252 | cell leading edge | 158/4138 | 7.52E-15 |
| GO:0005938 | cell cortex | 122/4138 | 9.92E-12 |
| GO:0030027 | lamellipodium | 85/4138 | 5.25E-11 |
| GO:0030139 | endocytic vesicle | 116/4138 | 3.43E-10 |
| GO:0010008 | endosome membrane | 164/4138 | 6.87E-10 |
| GO:0098589 | membrane region | 122/4138 | 7.21E-10 |
| GO:0098857 | membrane microdomain | 118/4138 | 1.06E-09 |
| GO:0045121 | membrane raft | 117/4138 | 1.74E-09 |
| GO:0005775 | vacuolar lumen | 70/4138 | 1.82E-07 |
| GO:0031983 | vesicle lumen | 114/4138 | 1.88E-07 |
| GO:0030864 | cortical actin cytoskeleton | 40/4138 | 8.69E-06 |
| GO:0001726 | ruffle | 63/4138 | 2.47E-05 |
| GO:0098858 | actin-based cell projection | 73/4138 | 2.47E-05 |
| GO:0005911 | cell-cell junction | 127/4138 | 3.61E-05 |
| GO:0030118 | clathrin coat | 21/4138 | 0.00108808 |
| GO:0019898 | extrinsic component of membrane | 88/4138 | 0.00137534 |
| GO:0001725 | stress fiber | 27/4138 | 0.001736 |
| GO:0097517 | contractile actin filament bundle | 27/4138 | 0.001736 |
| GO:0032432 | actin filament bundle | 29/4138 | 0.00232453 |
| GO:0005884 | actin filament | 39/4138 | 0.00254515 |
| GO:0032587 | ruffle membrane | 33/4138 | 0.006072 |
| GO:0044853 | plasma membrane raft | 37/4138 | 0.00632275 |
| GO:0070160 | tight junction | 42/4138 | 0.00651324 |

Supplementary Table 3. Gene Ontology: Molecular Functions

| ID | Description | GeneRatio | fDR |
|---|---|---|---|
| GO:0050839 | cell adhesion molecule binding | 200/4023 | 1.25E-15 |
| GO:0045296 | cadherin binding | 139/4023 | 1.60E-12 |
| GO:0003779 | actin binding | 164/4023 | 1.21E-10 |
| GO:0005096 | GTPase activator activity | 115/4023 | 1.57E-10 |
| GO:0030695 | GTPase regulator activity | 124/4023 | 2.40E-10 |
| GO:0017016 | Ras GTPase binding | 153/4023 | 7.85E-08 |
| GO:0031267 | small GTPase binding | 156/4023 | 1.21E-07 |
| GO:0051015 | actin filament binding | 83/4023 | 1.25E-07 |
| GO:0017048 | Rho GTPase binding | 76/4023 | 1.56E-07 |
| GO:0019902 | phosphatase binding | 77/4023 | 6.10E-07 |
| GO:0004674 | protein serine/threonine kinase activity | 151/4023 | 9.74E-07 |
| GO:0005178 | integrin binding | 54/4023 | 0.0001276 |
| GO:0016887 | ATPase activity | 138/4023 | 0.00034006 |
| GO:0005543 | phospholipid binding | 136/4023 | 0.0003452 |
| GO:0017049 | GTP-Rho binding | 14/4023 | 0.00086317 |

Supplementary Table 4. Signaling Pathway Impact Analysis: Significant Pathways

| Name | ID | pSize | NDE | pNDE | tA | pPERT | pG | pGFdr | pGFWER | Status | geneID |
|---|---|---|---|---|---|---|---|---|---|---|---|
| Focal adhesion | 4510 | 178 | 93 | 1.20E-12 | -49.5334 | 0.027 | 1.04E-12 | 1.42E-10 | 1.42E-10 | Inhibited | ITGB3/ITGB4/ITGB5/ITGA11/ITGA2/ITGA3/ITGA5/ITGAV/ITGA10/FLNA/FLNB/CDC42/MYLK/MYLK2/MYL12B/MYL9/PPP1CC/ROCK2/AKT3/PARVB/PARVA/ILK/PTEN/PIK3CD/PIK3CG/PIK3R3/TLN1/PIP5K1C/ACTN1/ACTN2/CAPN2/PXN/ARHGAP35/RHOA/COL1A1/COL1A2/COL4A1/COL4A2/COL4A6/COL5A1/COL5A2/COL6A1/COL6A3/FN1/LAMA2/LAMA5/LAMB1/LAMB2/LAMC1/LAMC2/RELN/SPP1/THBS1/THBS3/VTN/VWF/CAV1/CAV2/BIRC2/ELK1/MAPK1/MAPK3/CCND1/BRAF/RAF1/GRB2/EGFR/ERBB2/FLT4/IGF1R/KDR/MET/PDGFRA/IGF1/PGF/VEGFB/VEGFC/PDGFD/SHC4/SHC3/CRK/CRKL/RAPGEF1/DOCK1/JUN/PAK1/PAK2/RAC1/RAC2/RAC3/VAV3/CTNNB1/GSK3B |
| ECM-receptor interaction | 4512 | 72 | 41 | 1.06E-07 | -14.8166 | 0.031 | 6.76E-08 | 4.63E-06 | 9.26E-06 | Inhibited | LAMA2/LAMA5/LAMB1/LAMB2/LAMC1/LAMC2/COL1A1/COL1A2/COL4A1/COL4A2/COL4A6/COL5A1/COL5A2/COL6A1/COL6A3/GP6/VWF/VTN/SPP1/FN1/THBS1/THBS3/ITGB3/ITGAV/CD47/CD36/SV2A/SDC1/SDC4/SDC3/HSPG2/ITGB4/ITGB5/ITGA11/ITGA10/ITGA5/ITGA3/RELN/CD44/ITGA2/AGRN |
| Pathways in cancer | 5200 | 273 | 115 | 6.75E-08 | -32.1875 | 0.219 | 2.81E-07 | 1.28E-05 | 3.85E-05 | Inhibited | CTNNB1/WNT11/CTNNA1/COL4A1/COL4A2/COL4A6/FN1/LAMA2/LAMA5/LAMB1/LAMB2/LAMC1/LAMC2/ITGA2/ITGA3/ITGAV/PTEN/FZD5/BCR/CRK/CRKL/PIK3CD/PIK3CG/PIK3R3/STAT5B/JAK1/EGFR/ERBB2/PDGFRA/IGF1R/KIT/MET/FGFR1/TFG/TPM3/TPR/IGF1/HSP90AA1/HSP90AB1/HSP90B1/GSK3B/IKBKB/IKBKG/MTOR/MDM2/CDKN1B/BCL2L1/STAT3/STAT1/GRB2/RAF1/BRAF/RELA/CCND1/NCOA4/PLCG1/PLCG2/RALA/RALB/RHOA/RAC1/RAC2/RAC3/PLD1/CDC42/JUN/FOS/MYC/TP53/TCF7L1/STK36/FH/EGLN1/HIF1A/ARNT/EP300/MLH1/TGFB2/TGFBR1/TGFBR2/SMAD2/SMAD3/RUNX1/CTBP1/CTBP2/HDAC1/RXRA/JUP/RARA/PML/MITF/CDKN1A/CDK6/CCNE2/CDKN2A/BMP2/HHIP/BIRC2/TRAF5/PGF/VEGFB/VEGFC/SLC2A1/DAPK3/ETS1/MMP1/CXCL8/FOXO1/MAPK1/MAPK3/AKT3/GLI2/GLI3/KITLG/FGF14 |
| Pancreatic cancer | 5212 | 66 | 37 | 7.50E-07 | -19.2309 | 0.086 | 1.13E-06 | 3.88E-05 | 0.000155 | Inhibited | ERBB2/PIK3CD/PIK3CG/PIK3R3/ARHGEF6/AKT3/RAC1/RAC2/RAC3/IKBKB/IKBKG/RELA/BCL2L1/RAF1/BRAF/MAPK1/MAPK3/RALA/RALB/PLD1/JAK1/STAT3/STAT1/PGF/VEGFB/VEGFC/CDKN2A/CDC42/CDK6/TGFB2/TGFBR1/TGFBR2/SMAD2/SMAD3/TP53/EGFR/CCND1 |
| Regulation of actin cytoskele | 4810 | 177 | 74 | 1.90E-05 | -30.048 | 0.104 | 2.80E-05 | 0.000639 | 0.003832 | Inhibited | ARPC5/ARPC3/ARPC2/ARPC1A/PFN1/PFN2/BRK1/WASL/NCKAP1L/CYFIP1/CYFIP2/LIMK2/PIP5K1C/PIP4K2C/PIP5K1A/PIP4K2B/ACTN1/ACTN2/MSN/RDX/PXN/PPP1CC/MYL12B/MYL9/MYLK/MYLK2/GIT1/ENAH/ROCK2/PAK1/PAK2/ARHGEF6/CDC42/RAC1/RAC2/RAC3/RHOA/IQGAP3/ARHGAP35/VAV3/TIAM2/MAPK1/MAPK3/RAF1/BRAF/PIK3CD/PIK3CG/PIK3R3/DOCK1/CRK/CRKL/ITGA11/ITGA2/ITGA3/ITGA5/ITGAV/ITGAX/ITGB3/ITGB4/ITGB5/ITGA10/FN1/RRAS2/MRAS/RRAS/GNA12/EGFR/FGFR1/PDGFRA/FGF14/PDGFD/CD14/MYH9/MYH10 |
| MAPK signaling pathway | 4010 | 216 | 88 | 1.12E-05 | -8.35222 | 0.388 | 5.78E-05 | 0.00099 | 0.007918 | Inhibited | DUSP14/DUSP10/DUSP1/DUSP3/DUSP4/DUSP8/DUSP22/MAP4K4/PLA2G4A/PLA2G12A/RELA/RELB/IKBKB/IKBKG/MAPK1/MAPK3/RAF1/BRAF/NF1/PRKACB/PRKX/RRAS2/MRAS/RRAS/GNA12/GRB2/PDGFRA/FGFR1/EGFR/CACNG8/CACNG6/CACNA2D1/CACNB3/CACNA2D4/FGF14/BDNF/MAPK7/MAP2K5/NLK/CDC25B/RPS6KA4/MEF2C/TP53/ELK1/MAPK13/MAPK12/MAP2K6/AKT3/TAOK3/TAOK1/TAOK2/HSPB1/MAP3K5/MAP3K13/GADD45B/CD14/TGFBR1/TGFBR2/IL1R2/TNFRSF1A/TGFB2/MAPKAPK5/RAC1/RAC2/RAC3/CDC42/PAK1/PAK2/MAP3K3/MAPK8IP2/CRK/CRKL/ARRB1/JUND/JUN/NFATC4/NFATC2/CHP1/PPP3CA/PPP3CB/MAPK8IP3/FLNA/FLNB/MAP3K8/FOS/MYC/MKNK2/MKNK1 |
| Tight junction | 4530 | 102 | 48 | 1.46E-05 | -4.56989 | 0.375 | 7.17E-05 | 0.001091 | 0.009816 | Inhibited | F11R/JAM3/PRKCD/PRKCH/PPP2CA/OCLN/CLDN11/CLDN5/CLDN1/LLGL1/MAGI1/TJP1/EPB41/EPB41L1/EPB41L2/EPB41L3/CTNNA1/HCLS1/CSNK2A1/CSNK2B/SPTAN1/MYL12B/MYL9/MYH3/MYH9/MYH10/MYH7B/RHOA/ASH1L/EXOC4/PRKCI/RRAS2/MRAS/RRAS/AKT3/GNAI1/GNAI2/CTNNB1/CDC42/RAB13/ACTN1/ACTN2/CASK/TJP2/YES1/MAGI3/PTEN/AMOTL1 |
| VEGF signaling pathway | 4370 | 59 | 29 | 0.000275 | 16.54287 | 0.051 | 0.000171 | 0.001736 | 0.023401 | Activated | CDC42/KDR/PLCG1/PLCG2/PLA2G4A/PLA2G12A/HSPB1/CHP1/PPP3CA/PPP3CB/AKT3/PIK3CD/PIK3CG/PIK3R3/NFAT5/NFATC2/NFATC3/NFATC4/MAPK13/MAPK12/PXN/SH2D2A/MAPK1/MAPK3/RAF1/NOS3/RAC1/RAC2/RAC3 |
