## Supplementary figures and images for "AKAP12 variant 1 knockout enhances vascular endothelial cell motility"

### Supplemental original WB

**Suppl. Figure 1A original**

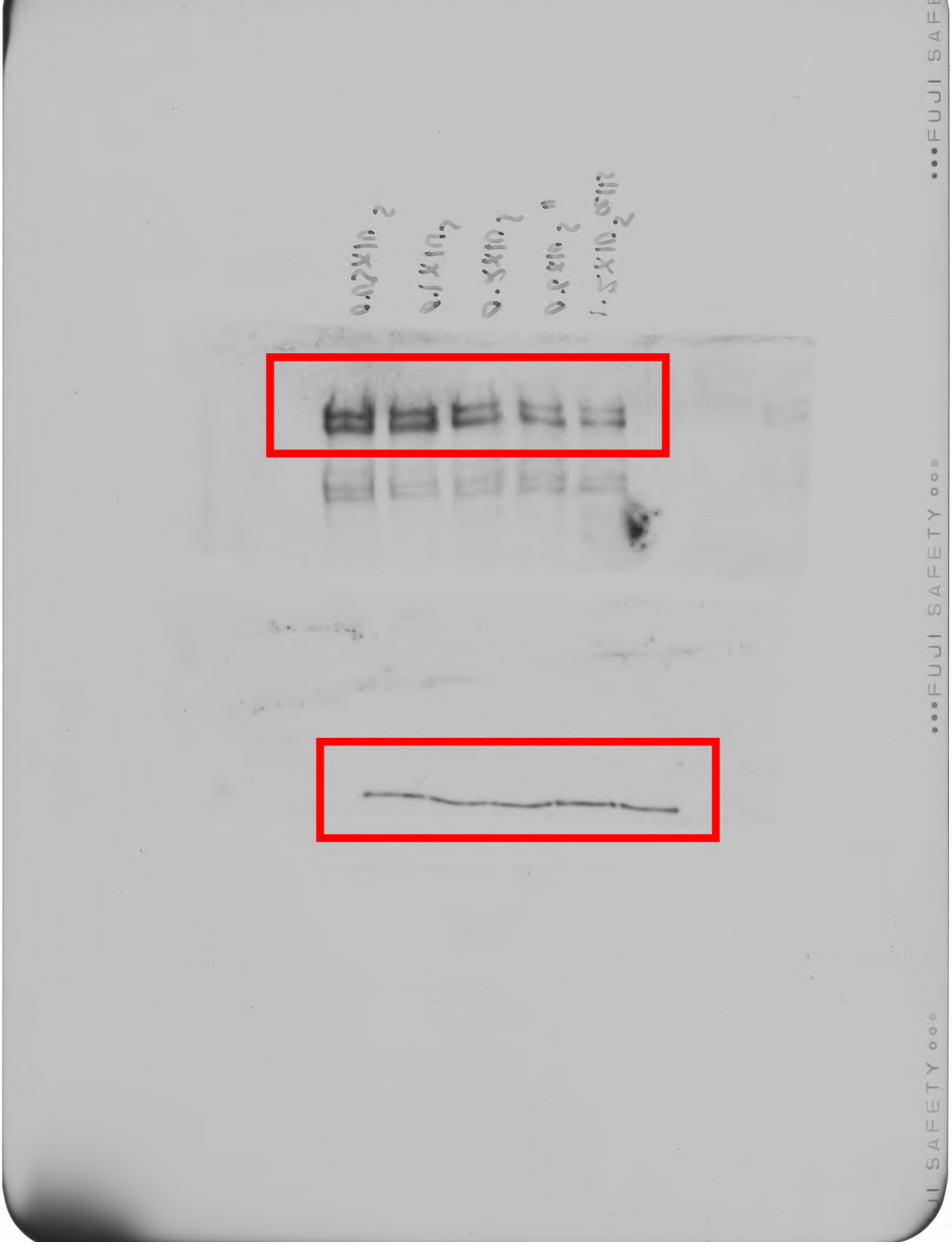

**Suppl. Figure 2A original**

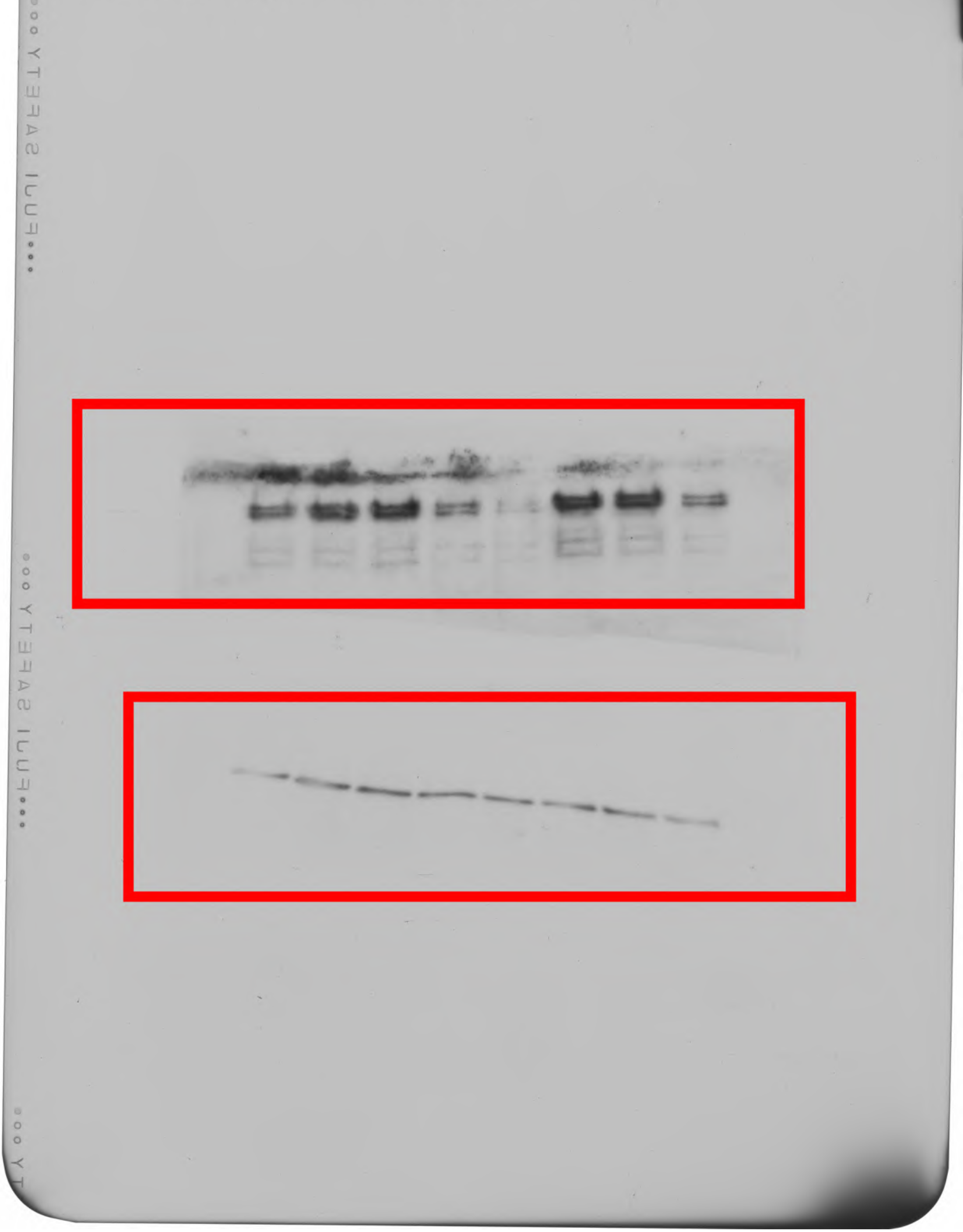

Suppl. Figure 3E original

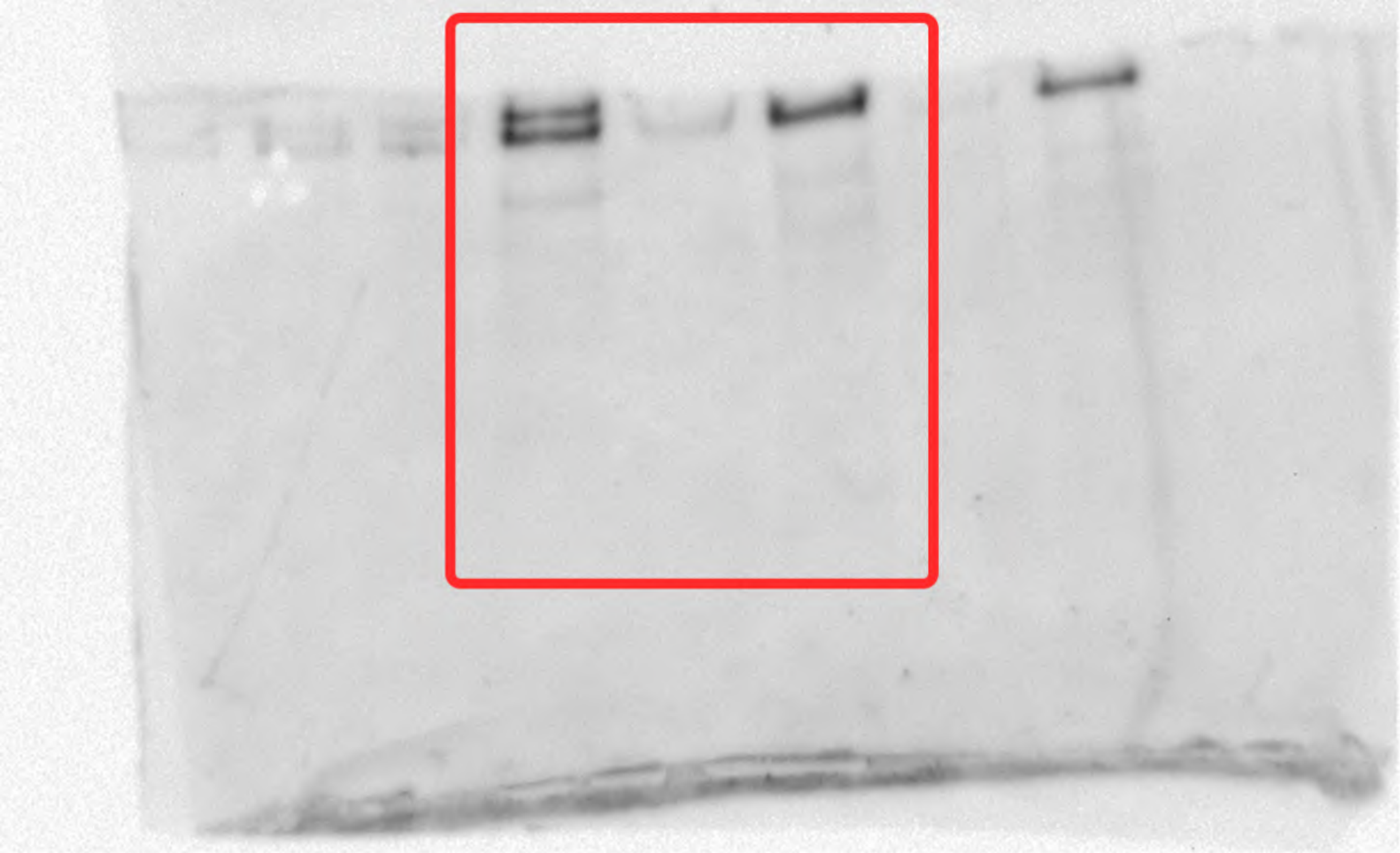

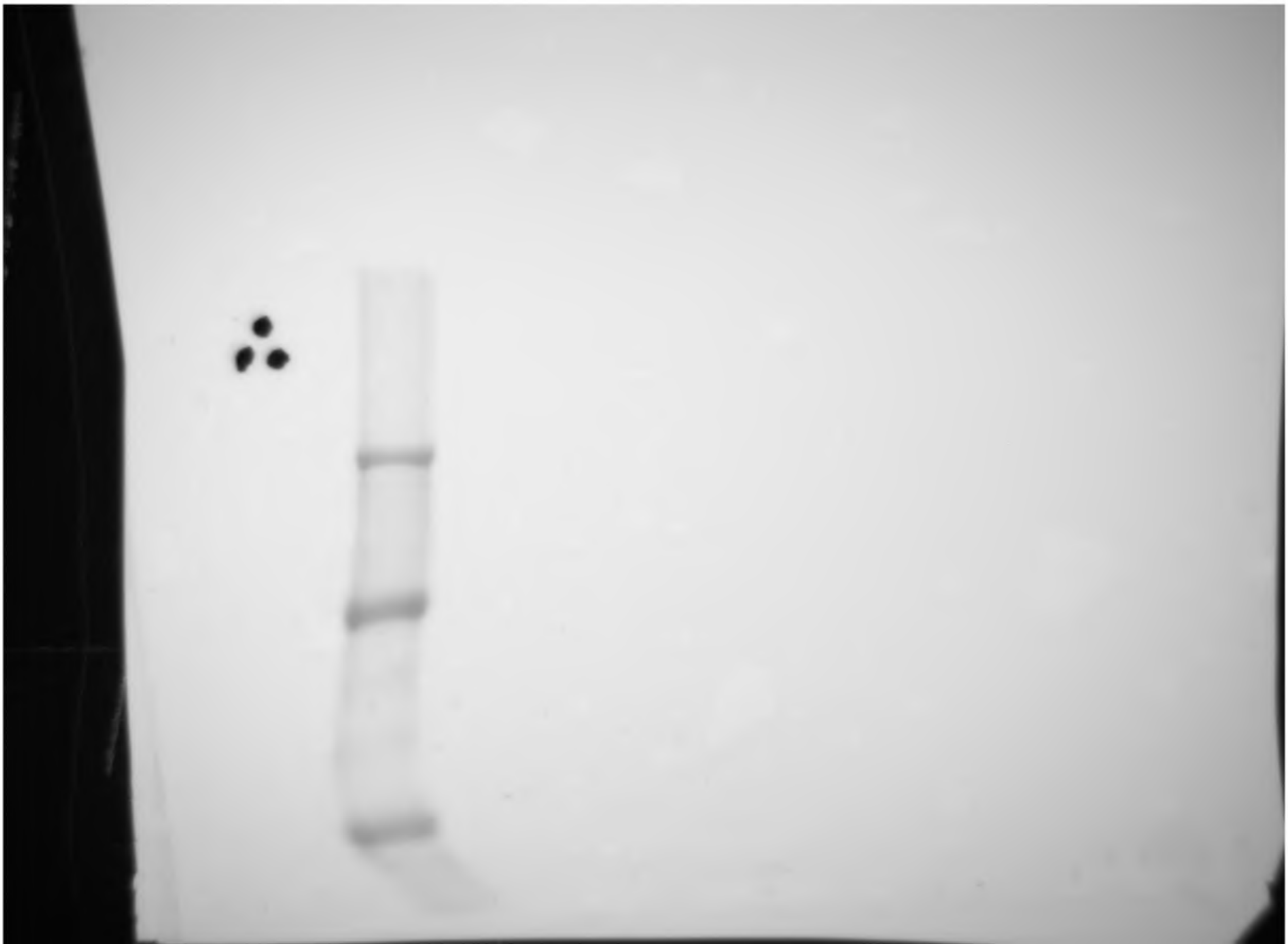
